# High-resolution analysis of the peptidoglycan composition in *Streptomyces coelicolor*

**DOI:** 10.1101/319178

**Authors:** Lizah T. van der Aart, Gerwin K. Spijksma, Amy Harms, Waldemar Vollmer, Thomas Hankemeier, Gilles P. van Wezel

## Abstract

The bacterial cell wall maintains cell shape and protects against bursting by the turgor. A major constituent of the cell wall is peptidoglycan (PG), which is continuously modified to allow cell growth and differentiation through the concerted activity of biosynthetic and hydrolytic enzymes. Streptomycetes are Gram-positive bacteria with a complex multicellular life style alternating between mycelial growth and the formation of reproductive spores. This involves cell-wall remodeling at apical sites of the hyphae during cell elongation and autolytic degradation of the vegetative mycelium during the onset of development and antibiotic production. Here, we show that there are distinct differences in the cross-linking and maturation of the PG between exponentially growing vegetative hyphae and the aerial hyphae that undergo sporulation. LC-MS/MS analysis identified over 80 different muropeptides, revealing that major PG hydrolysis takes place over the course of mycelial growth. Half of the dimers lack one of the disaccharide units in transition-phase cells, most likely due to autolytic activity. De-acetylation of MurNAc to MurN was particularly pronounced in spores, suggesting that MurN plays a role in spore development. Taken together, our work highlights dynamic and growth phase-dependent construction and remodeling of PG in *Streptomyces*.

**IMPORTANCE:** Streptomycetes are bacteria with a complex lifestyle, which are model organisms for bacterial multicellularity. From a single spore a large multigenomic, multicellular mycelium is formed, which differentiates to form spores. Programmed cell death is an important event during the onset of morphological differentiation. In this work we provide new insights into the changes in the peptidoglycan architecture over time, highlighting changes over the course of development and between growing mycelia and spores. This revealed dynamic changes in the peptidoglycan when the mycelia age, showing extensive PG hydrolysis and in particular an increase in the proportion of 3-3-cross-links. Additionally, we identified a muropeptide that is highly abundant specifically in spores, which may relate to dormancy and germination.

## INTRODUCTION

Peptidoglycan (PG) is a major component of the bacterial cell wall. It forms a physical boundary that maintains cell shape, protects cellular integrity against the osmotic pressure and acts as a scaffold for large protein assemblies and exopolymers (66). The cell wall is a highly dynamic macromolecule that is continuously constructed and deconstructed to allow for cell growth and to meet environmental demands (27). PG is built up of glycan strands of alternating N-acetylglucosamine (GlcNAc) and N-acetylmuramic acid (MurNAc) residues that are connected by short peptides to form a mesh-like polymer. PG biosynthesis starts with the synthesis of PG precursors by the Mur enzymes in the cytoplasm and cell membrane, resulting in lipid II precursor, undecaprenylpyrophosphoryl-MurNAc(GlcNAc)-pentapeptide. Lipid II is transported across the cell membrane by MurJ and/or FtsW/SEDS proteins and the PG is polymerized and incorporated into the existing cell wall by the activities of glycosyltransferases and transpeptidases (9, 30, 57).

The Gram-positive model bacterium *Bacillus subtilis* grows via lateral cell wall synthesis followed by binary fission; in addition, *B. subtilis* forms heat- and desiccation-resistant spores (23, 48). By contrast, the vegetative hyphae of the mycelial *Streptomyces* grow by extension of the hyphal apex and cell division results in connected compartments separated by cross-walls (2, 7, 11). This makes *Streptomyces* a model taxon for bacterial multicellularity (8). Multicellular vegetative growth poses different challenges to *Streptomyces*, including the synthesis of many chromosomes during vegetative growth and separation of the nucleoids in the large multi-genomic compartments during cross-wall formation (22, 68). Stress conditions such as nutrient depletion trigger the development of a so-called aerial mycelium which feeds on the underlying vegetative or substrate mycelium. The aerial hyphae eventually differentiate into chains of spores, a process whereby many spores are formed almost simultaneously, requiring highly complex coordination of nucleoid segregation and condensation and multiple cell division (22, 36, 41). Streptomycetes have an unusually complex cytoskeleton, which plays a role in polar growth and cell-wall stability (6, 19).

The *Streptomyces* genome encodes a large number of cell wall-modifying enzymes, such as cell wall hydrolases (autolysins), carboxypeptidases and penicillin-binding proteins (PBPs), a complexity that suggests strong heterogeneity of their PG (15, 45). Several concepts that were originally regarded as specific to eukaryotes also occur in bacteria, such as multicellularity (8, 31, 58), and programmed cell death (18, 51). Programmed cell death (PCD) likely plays a major role in the onset of morphological development, required to lyse part of the vegetative mycelium to provide the nutrients for the aerial hyphae (34, 38). PCD and cell-wall recycling are linked to antibiotic production in *Streptomyces* (33, 62).

All disaccharide peptide subunits (muropeptides) in the PG are variations on the basic building block present in lipid II, which in *Streptomyces* typically consists of GlcNAc-MurNAc-L-Ala-D-Glu-LL-DAP(Gly)-D-Ala-D-Ala (21, 56). Here, we have analyzed the cell wall composition of vegetative mycelium and mature spores of *Streptomyces coelicolor* by LC-MS, to obtain a detailed inventory of the monomers and dimers in the cell wall. This revealed extensive cell wall hydrolysis and remodeling during vegetative growth and highlights the difference in cell wall composition between vegetative hyphae and spores.

## RESULTS AND DISCUSSION

### PG isolation and analysis

*Streptomyces* form long hyphae and eventually large mycelial networks in submerged cultures (64). To assess how this translates to variations in the PG composition, we isolated the PG and analyzed the muropeptide profile of spores and of vegetative hyphae during different phases of growth in liquid-grown cultures. Vegetative mycelia of *S. coelicolor* were harvested from liquid minimal medium (NMM+). Samples taken after 18 and 24 h represented exponential growth, while samples taken after 36 h and 48 h represented mycelia in transition phase (Figure 1). Spores were harvested from SFM agar plates and filtered to exclude larger mycelial structures, all samples were taken in triplicate.

**Figure 1.**
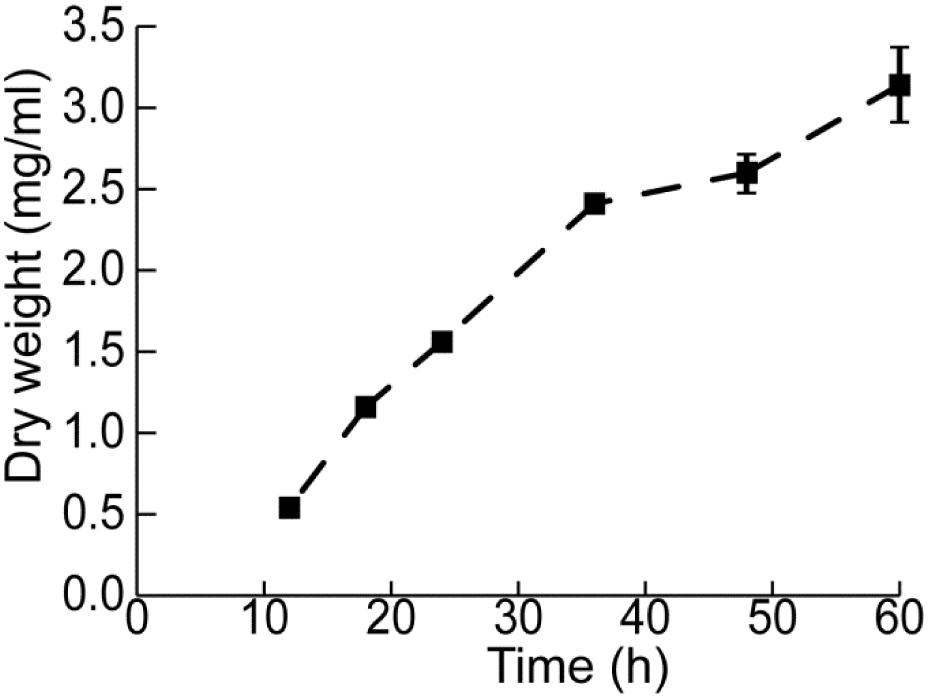
Growth of *S.coelicolor* on NMM+ medium based on triplicate dry weight measurements.

To allow analyzing a large number of samples simultaneously and in a reasonable time frame, we adapted a method for rapid PG purification (25) for *S. coelicolor*. For this, crude cell-wall material was isolated by boiling cells in 0.25% SDS and secondary cell wall polymers such as teichoic acids were removed by HCl. (see Materials and Methods section for details). As a control for the validity of the method, it was compared to a more elaborate method which is more routinely used (4), whereby *S. coelicolor* mycelia were treated with 5% boiling SDS and subsequently treated with hydrofluoric acid (HF) to remove teichoic acids. Comparison of the two methods revealed comparable outcomes between the two methods in peak detection (Table S3). Therefore, we proceeded with the method using 0.25%SDS and HCl.

The isolated PG was digested with mutanolysin (13, 25) and the muropeptide composition was analyzed by liquid chromatography linked to mass spectrometry (LC-MS). Every sample was analyzed three times and every sample in this dataset is run in the same sequence, to decrease possible deviations in retention time between the samples. Peaks were identified in the m/z range from 500-3000 Da, whereby different m/z in co-eluting peaks were further characterized by MS/MS. The eluted m/z values were compared to a dataset of theoretical masses of predicted muropeptides. Table 1 shows a summary of the monomers and dimers detected; the full datasets are given in Tables S1 and S2. We identified several modifications, including the amidation of D-iGlu to D-iGln at position 2 of the stem peptide, deacytelation of MurNAc to MurN, removal of amino acids to generate mono-, di-, tri- and tetrapeptides, loss of LL-DAP-bound glycine, and the presence of Gly (instead of Ala) at position 1,4 or 5. The loss of GlcNac or GlcNAc-MurNAc indicates hydrolysis (Figure 2).

**Figure 2.**
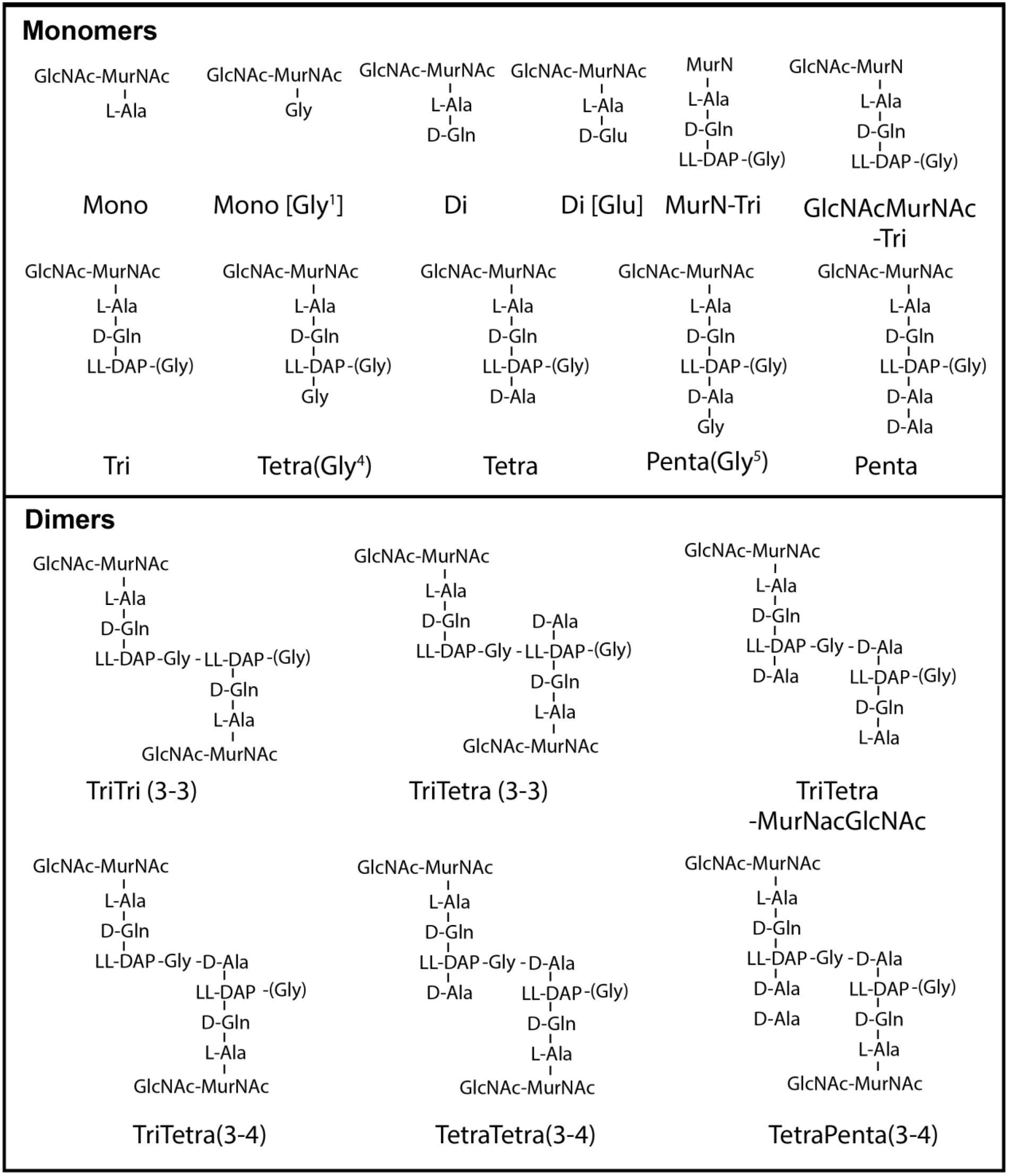
Summary of structures of main monomers and dimers observed in PG from *S. coelicolor*.

**Table 1.** Relative abundance(%) of muropeptides in vegetative cells and spores.

| **Monomers<sup>a</sup>** | **18h** | **24h** | **36h** | **48h** | **spores** |
| --- | --- | --- | --- | --- | --- |
| Mono | 1.6% | 2.1% | 3.3% | 3.3% | 4.5% |
| Di | 14.2% | 15.5% | 14.5% | 13.2% | 13.1% |
| Tri | 27.4% | 32.2% | 35.1% | 35.8% | 28.1% |
| Tetra | 26.7% | 24.4% | 23.9% | 23.9% | 48.3% |
| Tetra(Gly) | 3.5% | 5.3% | 6.9% | 8.2% | 2.3% |
| Penta | 22.7% | 16.9% | 13.1% | 12.9% | 5.3% |
| Penta(Gly) | 4.7% | 4.8% | 4.7% | 4.4% | 4.0% |
| Deacetylated | 3.9% | 6.0% | 7.9% | 8.0% | 4.5% |
| Glutamate | 67.6% | 63.4% | 62.6% | 66.3% | 74.0% |
| Missing GlcNAc | 0.7% | 1.6% | 2.1% | 3.3% | 4.8% |
| MurN-Tri | 0.1% | 0.7% | 1.2% | 2.3% | 3.5% |
| GlcNAcMurN-Tri | 2.2% | 2.8% | 3.4% | 2.6% | 0.2% |
| **Dimers<sup>a</sup>** | **18h** | **24h** | **36h** | **48h** | **spores** |
| TriTri (3-3) | 4.1% | 4.8% | 6.5% | 7.0% | 4.9% |
| TriTetra(3-3) | 23.9% | 24.2% | 22.3% | 16.9% | 19.1% |
| TriTetra(3-4) | 1.0% | 8.7% | 8.2% | 6.1% | 4.7% |
| TetraTetra(3-4) | 23.3% | 13.5% | 10.1% | 8.6% | 38.9% |
| TetraPenta (3-4) | 24.6% | 9.1% | 5.6% | 3.0% | 0.7% |
| Deacetylated | 1.8% | 1.2% | 1.5% | 1.2% | 0.4% |
| Glutamate | 66.5% | 61.1% | 62.8% | 62.0% | 70.5% |
| Missing GlcNAc | 0.3% | 0.6% | 1.1% | 1.2% | 0.1% |
| Missing (GlcNAc-MurNAc) | 24.3% | 37.2% | 44.6% | 56.1% | 30.4% |
| Proportion(%) of 3-3 cross-links | 36.5% | 48.0% | 54.5% | 57.3% | 35.1% |

<sup>a</sup>Monomers and dimers are treated as separate datasets.

### Fast growth rate correlates with high pentapeptide content

The first muropeptide after incorporation of Lipid II by glycosyltransferases contains a pentapeptide with a Gly residue linked to LL-DAP[3]. In many bacteria pentapeptides are short-lived muropeptides that occur mostly at sites of cell wall and cell division (26, 39). This is reflected by the high abundance of pentapeptides in the mycelia during exponential growth, with a pentapeptide content of 21% during early exponential growth (18 h) and 14% during late exponential growth (24 h) as compared to 11% in transition-phase cultures (36 h and 48 h). Conversely, tripeptides increased over time, from 24% during early exponential phase to 32% in transition-phase cultures.

### D-Ala is sometimes substituted for Gly

Gly can replace L-Ala[1] at position 1, or D-Ala[4/5] at positions 4 or 5 in in the pentapeptide. Addition of Gly to the medium and, in consequence, incorporation of Gly in the peptidoglycan can cause changes in morphology (16, 60) and facilitates lysozyme-mediated formation of protoplasts in *Streptomyces*, which was used to develop protoplast transformation methods (20, 24, 43). Gly at position 4 and 5 are incorporated in the murein, whether Gly at position 1 is incorporated at precursor level (60). Tetrapeptides carrying Gly at position 4 increased from 3% at 18 h, (exponential growth) to 8% at 48 h (transition and stationary phases) as the vegetative mycelium aged. The relative abundance of pentapeptides carrying Gly at position 5 was stable between 4-5% during growth.

### The abundance of 3-3 cross-links increases over time

It is possible to distinguish between two types of peptidoglycan cross-linking due to the difference in retention time and the application of MS/MS. These two types of cross-links are formed via two separate mechanism, the canonical D,D-transpeptidases (PBPs) produce a 3-4 (D,D) cross-links between LL-DAP[3] and D-Ala[4] and L,D-transpeptidases form 3-3 (L,D) cross-links between two LL-DAP[3] residues (Figure 2A). Dimers containing a tripeptide and a tetrapeptide (TetraTri) can have either cross-link, giving rise to isomeric forms that elute at different retention times, allowing for assessment by MS/MS (Figures 3A and 3B). We have found that the ratio of 3-3 cross-linking increased over time towards transition phase; the relative abundance increased from 37% of the total amount of dimers at 18 h (exponential phase) to 57% of all dimers at 48 h (Figure 2C).

**Figure 3.**
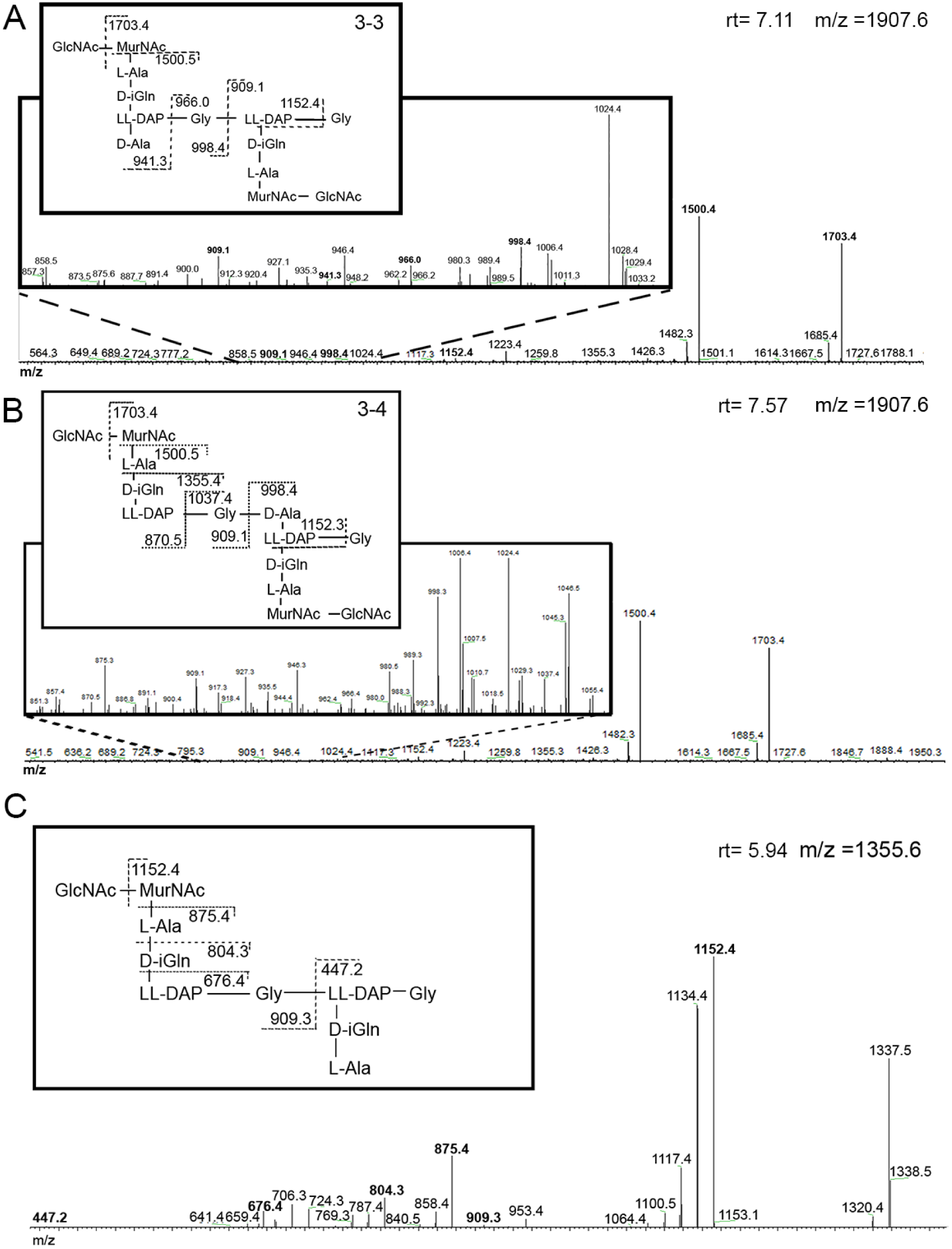
MS/MS fragmentations of TetraTri dimers with either 3-3 cross-link (A) or 3-4 cross-link (B). Differentiation between these two types of cross-links is possible at the point of asymmetry, at Gly attached to LL-DAP. The 3-3 cross-linked dimer (A) fragments into masses of 966.0 m/z and 941.3 m/z, which can be found in the respective MS/MS spectrum. The 3-4 cross-linked dimer (B) fragments into masses of 1037.4 m/z and 870.5 m/z. These masses are found in the MS/MS spectrum. Boxed MS/MS spectra show a magnification of masses between m/z 850 and 1050 to show masses present in lower abundance. (C) a TriTri dimer lacking GlcNAcMurNAc with an M+H of 1355.6, diagnostic fragments are given in the proposed structures.

### PG hydrolysis products increases as the culture ages

PG hydrolysis is associated with processes such as separation of daughter cells after cell division and autolysis, and in *E. coli* and other species deletion mutants lacking PG amidases grow in chains of unseparated cells (17, 67). Our data show that over the course of the growth curve, *S. coelicolor* increasingly lost GlcNAc and GlcNAc-MurNAc residues (Table 1); these are signs of cleavage by an N-acetylglucosamine and N-acetylmuramoyl-L-alanine amidase, respectively. The amount of dimers lacking GlcNAcMurNAc increases in time from 24% at 18 h to 56% at 48 h. Figure 3C shows MS/MS profiles of a Tri-Tri-dimer with a single set of glycans.

### Deacetylation of MurNAc is associated with mycelial aging

N-deacetylation of PG glycan strands is widespread among bacteria (65), which can occur both at GlcNAc and at MurNAc. Modifications to the glycan strand are commonly linked to lysozyme resistance (37). In the case of *S. coelicolor*, the only glycan modification is the deacetylation of MurNAc to MurN. Our data shows that this modification becomes more prominent as the vegetative mycelium ages, from 5% during early growth to 8% during later growth stages.

### MurN-Tri is prevalent in spores

The PG composition of spores and vegetative mycelia was compared to get more insights into the possible correlations between PG composition and important processes such as dormancy and germination. Our data show that muropeptides in spores are strongly biased for tetrapeptides, which make up 44% of the monomers, as compared to 23-25% of the vegetative PG. Conversely, pentapeptides were found in much lower amounts in spores at 5% of the monomers, whereas vegetative hyphae carry 10-22% of pentapeptides depending on the growth speed. A previous study has shown that a mutation in a D-alanyl-D-alanine carboxypeptidase (*dacA*) disrupts spore maturation and germination, where one could influence the other. This indicates that a either pentapeptides inhibit spore maturation, or that a high amount of tetrapeptides is essential (53). The muropeptide that stood out in the analysis of the spore PG was a tripeptide which lacks GlcNAc and contains a deacetylated MurNAc, called MurN-Tri. In spores, MurN-Tri made up 3.5% of the monomers, whereas the less modified muropeptide, GlcNAcMurN-Tri only made up 0.2% of the monomers. This suggests that MurN-tri is a spore-specific molecule. Glycan strand modifications are sometimes associated with germination site recognition (50), and we therefore speculate that the de-acetylated MurNAc residues could be recognized as germination sites of *Streptomyces* spores.

### Implications for the biology of actinobacteria

Sporulation is a key feature of *Streptomyces* biology and the spore wall is a major line of defense against environmental stresses, allowing the bacteria to survive under adverse conditions such as heat and cold stress, osmotic pressure, starvation or drought (44, 63). The spores are spread by e.g. insects or the wind, and germinate again in a new environment, as soon as the right conditions are met. Not much is known about the environmental and genetic factors that control the onset of germination. In terms of the genetics, we have previously shown that mutants deficient for the cAMP-receptor protein Crp have a much thicker spore wall, presumably due to reduced expression of cell-wall hydrolases and consequently germinate slowly (46). Additionally, Crp is an positive regulator of secondary metabolite production where the expression of Crp increases undecylprodigiosin production (12). Conversely, strains over-expressing the cell-division activator protein SsgA show an increase in the number of germ tubes per spore (42), with on average three germ tubes emerging from a single spore (instead of the two in wild-type spores and significantly less than that in *ssgA* mutants). It is yet unclear how future sites of branching in the hyphae or germination in the spores are marked. However, even after very long storage of spores, germination still occurs at the spore ‘poles’, suggesting physical marks to the PG, such as rare modifications. Yet, in our analysis, we did not find rare muropeptide moieties such as muramic δ-lactam, which is found in the spore-coat of *B. subtilis* (49). The only major difference was the relatively high amount of MurN-Tri in the spore PG (see above). It will be interesting to see why this moiety is overrepresented in the spore PG.

L,D-transpeptidases (LDTs) are especially prevalent in the actinobacterial genera *Mycobacterium*, *Corynebacterium* and *Streptomyces*, and, suggestively, these bacteria have a much higher percentage of 3-3 cross-links, with an abundance of at least 30% 3-3-cross links in investigated actinobacterial peptidoglycan. as compared to bacteria with lateral cell-wall growth such as *E. coli* (<10%) and *E. faecium* (3%) (5, 21, 28). LDTs attach to D-Ala[4] and form a cross-link between glycine and LL-DAP[3], D-Ala[5] is considered a donor for this type of cross-link (32). An interesting feature of these two mechanisms is that 3-4 cross-links can only be formed when a pentapeptide is present to display the D-Ala[5] donor, whereas 3-3 cross-links can be formed with a tetrapeptide as a donor strand. The ability to form 3-3 cross-links coincides with the ability to cross-link tetrapeptides instead of pentapeptides. We suggest that 3-3 cross-links could be essential to remodel the spore wall, using currently available muropeptides contrary to newly constructed pentapeptides (29, 54, 55).

### The cell wall and programmed cell death

Evidence is accumulating that, like eukaryotes, bacteria undergo a process of programmed cell death (PCD). Bacterial multicellularity implies a lifestyle involving cellular heterogeneity and the occasional sacrifice of selected cells for the benefit of survival of the colony (70–72). PCD is likely a major hallmark of multicellularity (8), and has been described in the biofilm-film forming *Streptococcus* (14) and *Bacillus* (10), in Myxobacteria that form fruiting bodies (59), in the filamentous cyanobacteria (3, 40), and in the branching *Streptomyces* (35, 38, 61). In streptomycetes, cell-wall hydrolases support developmental processes like branching and germination (15). Additionally, PCD is likely an important event during the onset of development from a vegetative lifestyle to a reproductive lifestyle, as autolytic degradation of the cell wall is intrinsically linked with the onset of antibiotic production and spore formation (61). Conceptually, GlcNAc accumulates at high concentrations around colonies during PCD, and GlcNAc is an important signaling molecule for the onset of morphological differentiation and antibiotic production in streptomycetes (52, 62). The linkage of PCD and antibiotic production is logical from a biological perspective; autolytic dismantling of the vegetative (substrate) mycelium generates building blocks in a nutrient-depleted soil, which will inevitably attract motile competitors. Antibiotics likely serve to fend off these competitors and protect the food source. Thus, cell-wall hydrolysis may facilitate the correct timing of development. As shown in this work, *Streptomyces coelicolor* carries ’scars’ of previous cell-wall hydrolysis events during the entire lifecycle, which emphasizes the importance of such hydrolysis to support cell growth.

## CONCLUSIONS

We have provided a detailed analysis of the peptidoglycan of *Streptomyces* mycelia and spores, and developed a reliable and fast method to compare larger numbers of samples. Our data show significant changes over time, among which changes in the amino acid chain, hydrolysis of dimers, and the accumulation of the rare MurN-Tri specifically in the spores. The cell wall likely plays a major role in the development of streptomycetes, with implications for germination and the switch to development and antibiotic production (via PCD-released cell wall components). The dynamic process that controls the remodeling of the cell wall during tip growth is poorly understood, but we anticipate that the local cell-wall structure at sites of growth and branching may well be different from that in older (non-growing) hyphae. This is consistent with the changes we observed over time, between the younger and older mycelia. Detailed localization of cell-wall modifying enzymes and of specific cell-wall modifications, in both time and space, is required to further reveal the role of the cell wall in the control of growth and development of streptomycetes.

## EXPERIMENTAL PROCEDURES

### Bacterial strain and culturing conditions

*Streptomyces coelicolor* A3(2) M145 (24) was obtained from the John Innes Centre strain collection. All media and methods for handling *Streptomyces* are described in the *Streptomyces* laboratory manual (24). Spores were collected from Soy Flour Mannitol (SFM) agar plates. Liquid cultures were grown shaking at 30°C in a flask with a spring, using normal minimal medium with phosphate (NMM+) supplemented with 1% (w/v) mannitol as the sole carbon source; polyethylene glycol (PEG) was omitted to avoid interference with the MS identification. Cultures were inoculated with spores at a density of 10^6^ CFU/ML. A growth curve was constructed from dry-weight measurements by freeze-drying washed biomass obtained from 10 mL of culture broth (three biological replicates). The respective aliquots were immediately imaged to show the pellet morphology. Spores were collected from SFM agar plates by adding 0.01% (w/v) SDS to facilitate spore release from the aerial mycelium, scraping them off with a cotton ball and drawing the solution with a syringe. Spores were filtered with a cotton filter to separate spores from residual mycelium.

### PG extraction

Cells were lyophilized for a biomass measurement, 10 mg biomass was directly used for PG isolation. PG was isolated according to (25), using 2 mL screw-cap tubes for the entire isolation. Biomass was first boiled in 0.25% SDS in 0.1 M Tris/HCl pH 6.8, thoroughly washed, sonicated, treated with DNase, RNase and trypsin, inactivation of proteins by boiling and washing with water. Wall teichoic acids were removed with 1 M HCl. PG was digested with mutanolysin and lysozyme (1). Muropeptides were reduced with sodium borohydride and the pH was adjusted to 3.5-4.5 with phosphoric acid.

To validate the method, we compared it to the method described previously (4). For this, *S. coelicolor* mycelia were grown in 1 L NMM+ media for 24 h. After washing of the mycelia, pellets were resuspended in boiling 5% (w/v) SDS and stirred vigorously for 20 min. Instead of sonicating the cells, they were disrupted using glass beads, followed by removal of the teichoic acids with an HF treatment at 4°C as described.

### LC-MS analysis of monomers

The LC-MS setup consisted of a Waters Acquity UPLC system (Waters, Milford, MA, USA) and a LTQ Orbitrap XL Hybrid Ion Trap-Orbitrap Mass Spectrometer (Thermo Fisher Scientific, Waltham, MA, USA) equipped with an Ion Max electrospray source. Chromatographic separation was performed on an Acquity UPLC HSS T3 C_18_ column (1.8 μm, 100 Å, 2.1 × 100 mm). Mobile phase A consist of 99.9% H_2_O and 0,1% Formic Acid and mobile phase B consists of 95% Acetonitrile, 4.9% H_2_O and 0,1% Formic Acid. All solvents used were of LC-MS grade or better. The flow rate was set to 0.5 ml/min. The binary gradient program consisted of 1 min 98% A, 12 min from 98% A to 85% A, and 2 min from 85% A to 0% A. The column was then flushed for 3 min with 100% B, the gradient was then set to 98% A and the column was equilibrated for 8 min. The column temperature was set to 30°C and the injection volume used was 5 μL. The temperature of the autosampler tray was set to 8°C. Samples were run in triplicates.

MS/MS was done both on the full chromatogram by data dependent MS/MS and on specific peaks by selecting the mass of interest. Data dependent acquisition was performed on the most intense detected peaks, the activation type was Collision Induced Dissociation (CID). Selected MS/MS was performed when the resolution of a data dependent acquisition lacked decisive information. MS/MS experiments in the ion trap were carried out with relative collision energy of 35% and the trapping of product ions were carried out with a q-value of 0.25, and the product ions were analyzed in the ion trap., data was collected in the positive ESI mode with a scan range of *m*/*z* 500–3000 in high range mode. The resolution was set to 15.000 (at m/z 400).

### Data analysis

Chromatograms were evaluated using the free software package MZmine (http://mzmine.sourceforge.net/ (47)) to detect peaks, deconvolute the data and align the peaks. Only peaks corresponding with a mass corresponding to a muropeptide were saved, other data was discarded. The online tool MetaboAnalyst (69) was used to normalize the data by the sum of the total peak areas, then normalize the data by log transformation. The normalized peak areas were exported and a final table which shows peak areas as percentage of the whole was produced in Microsoft Excel.

### Muropeptide identification

The basic structure of the peptidoglycan of *S. coelicolor* has been published previously (21). Combinations of modifications were predicted and the masses were calculated using ChemDraw Professional (PerkinElmer). When a major peak had an unexpected mass, MS/MS helped resolve the structure. MS/MS was used to identify differences in cross-linking and to confirm predicted structures.

## Acknowledgments

This work is part of the profile area Antibiotics of the Faculty of Sciences of Leiden University.

## Author contributions

LvdA performed the experiments with the help of GS. LvdA and GvW conceived the study. LvdA, AH, TH and GvW wrote the article with the help of WV. All authors approved the final manuscript.

